# The migration game in habitat network: the case of tuna

**DOI:** 10.1101/020743

**Authors:** Patrizio Mariani, Vlastimil Křivan, Brian R MacKenzie, Christian Mullon

## Abstract

Long distance migration is a widespread process evolved independently in several animal groups in terrestrial and marine ecosystems. Many factors contribute to the migration process and of primary importance are intra-specific competition and seasonality in the resource distribution. Adaptive migration in direction of increasing fitness should leads to the ideal free distribution (IFD) which is the evolutionary stable strategy of the habitat selection game. We introduce a migration game which focuses on migrating dynamics that lead to the IFD for age-structured populations in time varying environments where dispersal is costly. The model assumes a network of habitats and predicts migration dynamics between these habitats and the corresponding population distribution. When applied to Atlantic bluefin tunas it predicts their migration routes and their seasonal distribution. The largest biomass is located in the spawning areas which have also the largest diversity in the age-structure. Distant feeding areas are occupied on a seasonal base and often by larger individuals, in agreement with empirical observation. Moreover we show that only a selected number of migratory routes emerge as those effectively used by tunas.

## 1 Introduction

Many populations of animals and plants exhibit characteristic distributional patterns that are related to the ability of the organisms to move and explore their environment. Changes in the environment can elicit individual reactions, hence causing different spatial distributions of populations (Morris, 2011). Competition is among the major driving forces shaping animal distributions. Dispersal from more populated to less populated habitats reduces intra-and inter-specific competition thus promoting species coexistence and diversity (MacArthur and Levins, 1964; Rosenzweig, 1981). These dispersal dynamics often involve active habitat selection, which is a widespread phenomenon in nature and has been described in many animal populations such as birds (Cody, 1985), terrestrial mammals (Wecker, 1963) and fish (MacCall, 1990).

Fitness based arguments are commonly used to describe of the process of habitat choice (MacArthur and Levins, 1964). When moving between different habitats, organisms should prefer those sites that provide them with the highest payoff, i.e., where their fitness is maximised (Rosenzweig, 1981). Nevertheless, both individual fitness and habitat selection typically depend on interactions between individuals, which usually have the form of a density dependent relation linking habitat quality and species distribution (Rosenzweig and Abramsky, 1985).

Under negative density dependence (described by logistic growth), if dispersal is cost free and individuals are omniscient and free to settle at any habitat, the evolutionarily stable strategy corresponds to the ideal free distribution (IFD) (Fretwell and Lucas, 1969; Křivan et al, 2008; 2011). At the IFD, payoffs in all occupied habitats are the same and larger or equal than those in the unoccupied habitats. Thus, no individual can improve its fitness by choosing a different habitat.

Although the IFD is a strong theoretical tool to analyse animals’ spatial distributions, over the past decades attempts to validate it led to equivocal results (Matsumura et al, 2010). For example, several studies reported “under matching” when animals underuse better patches and overuse poorer patches (for a review see Kennedy and Gray, 1993). These discrepancies between the IFD and observed distributions are attributed to, e.g., the cost of moving (Åström, 1994), imperfect information (Matsumura et al, 2010), or stochastic fluctuations in environmental conditions (Schreiber, 2012).

An important aspect that is usually neglected in theoretical studies of habitat selection and migration (but see, e.g., Sutherland and Parker, 1985; Hugie and Grand, 1998; Grand and Dill, 1999; Tregenza and Thompson, 1998) is the variability among individuals. In particular, factors related to age or energetic state can contribute to individuals’ perception of the environment and affect the ability to migrate between habitats. Moreover, the specific location in which each individual lives can also affect the habitat selection process. Indeed, while the IFD assumes freely moving individuals between habitats, habitat-connectivity can often be constrained by specific geographical (e.g., topography) or temporal (e.g., seasonal) patterns, which can then limit the ability to migrate towards better habitats. A network of habitats is often a more realistic and general description of habitat connectivity for migratory species.

For example, migratory species such as the Atlantic bluefin tuna (BFT) have widely separated feeding and spawning areas that are distributed over a large latitudinal gradient. Those habitats are typically exposed to changes in seasonality and habitat productivity that can affect payoffs and dispersal dynamics of the habitat selection game. The species appears to have evolved a migration strategy that alternates rapid movement between neighbouring regions, to periods of continuous feeding in those areas before a new migration occurs (Block et al, 2005, 2001; Wilson et al, 2005). Thus, the dispersal dynamic between distant habitats appears as a multiple step process by which tunas explore several habitats rather than a single direct movement towards higher payoff areas.

In this manuscript we present a game theoretical approach, called the “migration game”, to model migration dynamics of an age-structured population on a network of interconnecting habitats that undergo seasonal variation. In addition, we assume a travel cost that is age specific. Then we apply this concept to BFT to predict their seasonal distribution and their migration routes across the Atlantic.

## 2 Theoretical framework

### 2.1 The migration game

We consider an unstructured migratory species in an heterogeneous environment consisting of a network with *n* habitats. Population and distributional processes are assumed to be discrete in time, and the time step is scaled so that it equals 1. In each habitat, *i*, and at each time step the population abundance, *p*_*i*_, changes due to migration and population dynamics:

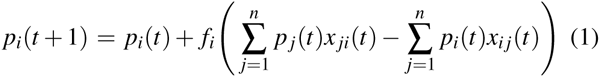

where *f*_*i*_ is the demographic change of the population *pi* (birth and death processes), and *x_ij_*(*t*) is the per capita migration rate from patch *i* to patch *j* within the unit time interval. The model (Eq. 1) assumes that in each time interval dispersal occurs before demographic changes. The total population abundance at time *t* is 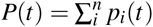. Dispersal rates, *x_ij_*(*t*), are non-negative and satisfy 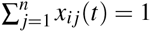 for every *i* = 1,…,*n*. We note that *x_ii_*(*t*) is the probability of staying in the patch *i*.

To describe migration rates we assume that each habitat is characterised by a negative density dependent payoff, *u*_*i*_. If there is a direct link between habitats *i* and *j* in the network, then for individuals migrating from *i* to *j* we define a reward function:

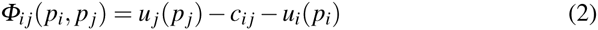

where *c_ij_* ≥ 0 is a cost term for the migration game. This cost includes the energy needed to migrate between habitats *i* and *j* as well as the energy required for habitat selection and decision making processes (Bonte et al, 2012).

We consider directed (non-random) movements on the network and we assume that along the migration routes the reward must be non-negative. In other words, an individual currently in patch *i* will move to patch *j* only when the reward of doing so is positive. Hence at each time step, *t*, dispersal rates *x_ij_* must results in a population distribution satisfying:

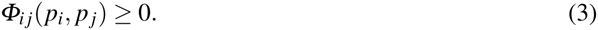

In the model motility is restricted by the topology of the network and individuals can only migrate between neighbouring habitats (i.e., habitats directly connected by a link). Moreover, the choice of an individual affects migration rates, *x_ij_*, and also population distribution (*pi*, Eq. 1). Hence, the rewards (*Φ*_*ij*_, Eq. 2) are regulated by the reciprocal strategies of competing individuals.

This defines a non cooperative migration game, in which (1) players are the set of individuals characterised by their current habitat *i*, (2) the strategy of the players currently in habitat *i* is the probability *xij* with which the individuals move to one of the neighbouring habitats, and (3) the reward of the set of players living in *i* is defined as the average reward ∑_*j*_ *x*_*ij*_*Φ*_*ij*_ where the sum is only over all neighbouring habitats *j* (i.e., habitats directly connected to habitat *i*).

The equilibrium solutions are migration rates 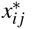 that are the Nash equilibria (NE) of the migration game. The equilibrium strategy is such that any unilateral change in the strategy of any individual would results in a lower reward for the player who changes its strategy. This implies that for any two habitats *j* and *j′* such that 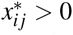 and 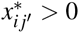, the rewards must be the same and maximal (i.e., 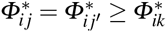 for any connected habitat *k* such that *x_ik_* = 0). The migration game is a potential game that guarantees the existence and uniqueness of a NE (Sandholm, 2010). This equilibrium can be calculated as the solution of a variational inequality (Nagurney, 1993; Mullon and Nagurney, 2012; Nagurney et al, 1992) or a linear complementary problem (LCP) (Mullon, 2013; Facchinei and Pang, 2003).

Migration rates 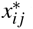 are then used in the model (Eq. 1) to define population dynamics on the network.

### 2.2 Distributional equilibrium in a cost-free migration game

We assume that migration is cost free, i.e., *c_ij_* = 0 in Eq. (2), hence the reward function is similar to those used in habitat selection games (Hugie and Dill, 1994; Křivan et al, 2008) and the solution of the migration game converges in several time steps to the IFD (Pan and Nagurney, 1994; Cressman and Křivan, 2006). However, because dispersing animals are constrained in their movement by links in the habitat network, depending on the topology of the network it can take several steps to reach the global IFD. Indeed at each time step individuals can move only to habitats that are directly connected to their current habitat. Thus, at each time step individuals reach a local IFD in the sense that directly linked patches have the same payoffs as we do not consider the cost of dispersal. As time increases, the IFD becomes more global, that is, payoffs in additional patches get equalised.

To illustrate the relation between migration equilibrium and distributional equilibrium we consider a simple case of three habitats denoted as *A*, *B*, *C*, and two different network topologies: (a) a fully connected network (Figure 1a); (b) a network where B is disconnected from C (Figure 1b). Each habitat is characterised by its payoff:

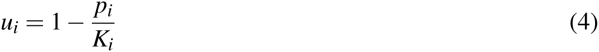

where *K*_*i*_ is the habitat environmental carrying capacity, *p*_*i*_ is the number of individuals in patch *i* with *i* ∈ {*A*,*B*,*C*}. We assume no migration costs (*c_ij_* = 0) and all individuals, *P*, initially occupying habitat *C* only, i.e., *p*_*C*_ = *P*, *p*_*A*_ = *p*_*B*_ = 0.

**Fig. 1.**
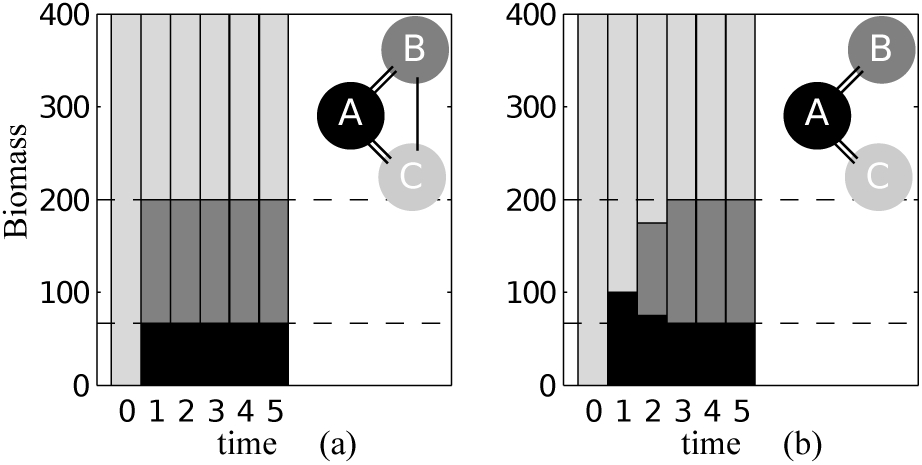
Cumulative distribution of the migratory population in the three habitats A, B, C in the case (a) all the habitats are connected (b) connections between A = B and A = C. Also shown (dashed lines) the reference cumulative distribution at the IFD. Parameters: *K*_*A*_ = 100, *K*_*B*_ = 200, *K*_*C*_ = 300 and total population *P* = 400.

When the network is fully connected our model converges to the IFD in a single time step (Figure 1a). Since individuals are free to move in all the habitats in the network, the strategies resulting from the migration game are those needed to balance the reward function in Eq. (2) for all the three habitats. This condition is also the condition for the IFD.

When the network is not fully connected, several steps are needed to reach the IFD (Figure 1b). The equilibrium value can be efficiently calculated using variational inequality (Mullon, 2013). In the first step, only movements between *C* and *A* are possible on the network and the values of *x_ij_* are those balancing the rewards (*Φ*_*CA*_ = *Φ*_*AC*_), i.e., a local IFD conditions is reached between the two habitats. In the next step, individuals that are now in habitat *A* have the possibility to migrate into *B* since *ΦAB* > 0. But, not all the dispersal rates are possible to reach the equilibrium, since for any given habitat the number of migrants cannot be larger than the number of inhabitants. Indeed, the migration equilibrium *x_AB_*, *x_CA_* must satisfy constraints 0 ≤ *x_AB_* ≤ *p*_*A*_ and 0 ≤ *x_CA_* ≤ *p*_*C*_. The new distribution is again calculated equalising the reward functions in the three habitats and considering that after the migration the payoff in habitat *A* can be written as

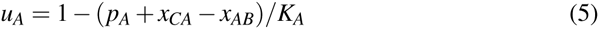

with similar expressions for the payoff in habitat *B* and *C*. Equilibrium values for *x_CA_* and *x_AB_* are calculated by equalising the patch payoffs. In particular, at this step all the individuals living in *A* move into *B* and a new distribution is reached between *A* and *C*. At step number three, the equilibrium values for dispersal rates are those satisfying the IFD on the network (Figure 1b).

### 2.3 The effects of costs and multiple equilibria

When travel costs are zero and patch payoffs are negative density dependent there is a single IFD (Křivan et al, 2008). However, if migration costs are positive there may be infinitely many possible IFDs. Indeed, let us consider an environment consisting of two habitats (*i* = 1,2), and let the habitat payoffs be described by the Eq. (4).

The reward of an individual currently in patch 1 to migrate to patch 2 is

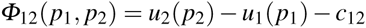

and, similarly, the reward of an individual currently in patch 2 to migrate to patch 1 is

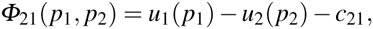

Under the IFD none of these two rewards can be positive. In particular, when travel costs are neglected a single IFD exists at which *Φ*_12_(*p*1, *p*2)= *Φ*21(*p*1, *p*2)= 0 (Figure 2A). When travel costs are positive (Figure 2B) there is a region of possible distributions under which individuals in neither patch have a positive reward to move. Thus, all these distributions correspond to IFDs.

**Fig. 2.**
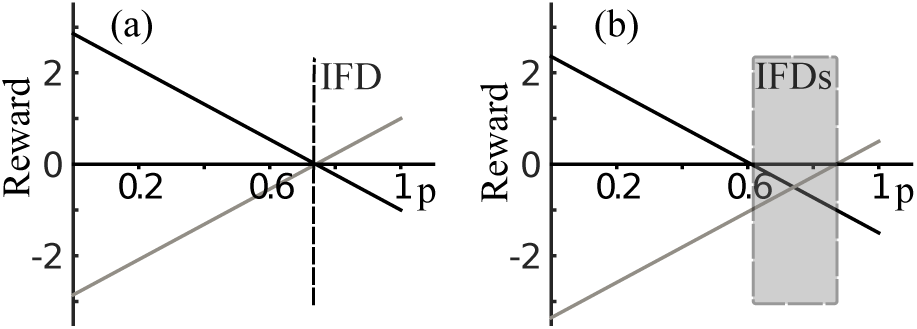
Reward functions (*Φ*_*ij*_) with and without migration costs. The plots show rewards of migration for individuals in patch 1 (red) and in patch 2 (blue) as the proportion *p* of individuals in patch 1. For values of *p* such that the red (blue) line is above the x-axis, individuals of population 1 (2) have advantage to migrate. In (a) migration costs are zero and a single equilibrium is present at *p* = 0.74. While in (b) migration costs are positive and there is a set of equilibrium distributions (*p* ∈ [0.58,0.84]).

The LCP method that we use to calculate numerically the NE of the migration game selects a single IFD from the set of possible IFDs, The selected point is on the boundary of the set of all IFDs, i.e., in the above example it is one of the two boundary points.

### 2.4 Coupling migration and demographic processes

We extend the model (Eq. 1, 2) by considering a structured population with several (*S*) age classes *a*. It can be represented as 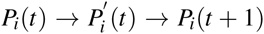, where *P*_*i*_(*t*)= {*p*_*i*,*a*_} is the population distribution, once individuals have redistributed themselves according to the migration equilibrium, and *P*_*i*_(*t* + 1) is the population distribution after the demographic processes (birth, death and growth). To represent these processes, in each habitat, we use a Leslie matrix, i.e., we have 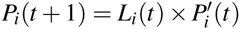 where:

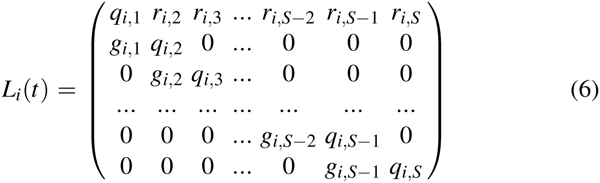

The probability that a fish in class *a* − 1 at time *t* will grow into class *a* at time *t* +1 is *g*_*a*_. Similarly, *q*_*a*_ is the probability that a fish will continue to stay in the same class, while *r*_*a*_ is the average number of newborns (belonging to the class *a* = 1) produced by individuals at ages *a* > 1.

## 3 Case Study

### 3.1 The ecology of Atlantic bluefin tuna

The Atlantic bluefin tuna (*Thunnus thynnus*) has evolved a migratory behaviour in which spawning and feeding sites are separated by large distances, typically spanning 100s1000s of kilometres and several degrees of latitude (Mather et al, 1995; Cury et al, 1998). Spawning sites are located in temperate-tropical waters (i.e., Mediterranean Sea, Gulf of Mexico), but feeding sites used by the largest and oldest individuals are located in northern temperate-boreal waters (Mather et al, 1995). During the narrow reproductive period individuals often display fast trans-Atlantic migrations to reach the Mediterranean spawning ground (Block et al, 2005; Fromentin, 2009). The seasonal south-north migratory behaviour exhibited by bluefin tuna has likely evolved to allow the species to benefit from large biomasses of prey species in these regions (Cury et al, 1998).

### 3.2 Model implementation

We implement the theoretical framework described above, to illustrate the spatial dynamics of the Atlantic Bluefin tuna. The time step for the dynamic system is set equal to one month, and the simulations are extended up to 20 years. We chose a network of *n* = 8 habitats (Figure 3): Gulf of Mexico (A), Brazil (B), Maine (C), North Atlantic (D), Norway (E), Bay of Biscay (F), eastern Atlantic (G) and Mediterranean (H). The links between habitats are selected based on historical migration routes of bluefin tuna and defined to represent feasible distances that individuals can cover in one month. Moreover, the migratory population is structured in five classes (age, *a*): young of the year, juvenile, adult, mature and old. We denote by *w*_*a*_ the mean weight of tuna at age class *a* (Table 1).

**Fig. 3.**
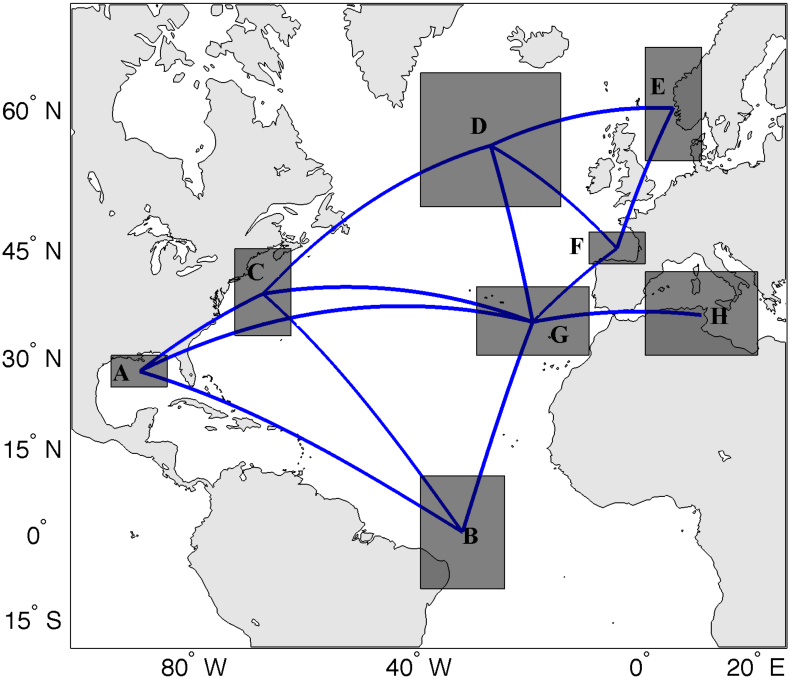
Network for the bluefin tuna migration game. We consider 8 habitats: A: Gulf of Mexico, B: Brazil, C: Maine, D: North Atlantic, E: Norway, F: Bay of Biscay, G: Eastern Atlantic, H: Mediterranean. Habitats are defined within a certain spatial range (grey areas) for which we calculate the average biological productivity which is assumed proportional to the habitat’s carrying capacity. Links between habitats indicates the potential migration routes assumed in the present study.

**Table 1.** Biological characteristics of age classes

| Age class<br>$a$ | Weight<br>$w$ (Kg) | Fertility<br>$r^a$ ($month^{-1}$) | Growth<br>$g$ ($month^{-1}$) | Survival<br>$q$ ($month^{-1}$) |
| --- | --- | --- | --- | --- |
| Young | 1 | 0 | 0.02 | 0.9 |
| Juvenile | 30 | $0.125s$ | 0.02 | 0.9 |
| Adult | 100 | $0.25s$ | 0.02 | 0.9 |
| Mature | 200 | $0.5s$ | 0.02 | 0.9 |
| Old | 500 | $s$ | 0.02 | 0.9 |

Patch payoffs for an individual of class *a* in habitat *i* ∈ {*A*,…,*H*} at month *t* are density dependent:

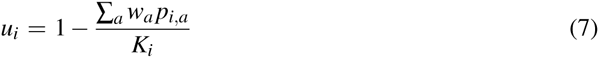

where *p*_i,*a*_ is the population of age *a* living in habitat *i*; *K*_*i*_ is the time varying carrying capacity of habitat *i* described as *K*_*i*_(*t*)= *K*_*i*_(1 − *θ*_*i*_ cos(*tπ*/2)), with *θ_i_* ≤ 1, being a seasonality parameter specific for each habitat (Table 2). The seasonality parameter is calibrated using averaged data (2003 -2011) of seasonal variability of the biological productivity (Westberry et al, 2008) averaged over the area covered by the habitat (Figure 3). Larger coefficients reflect larger seasonal fluctuations typically at higher latitudes.

**Table 2.** Characteristics of habitats

| Habitat $i$ | Mean carrying capacity $K_i$ | Seasonal effect $\theta_i$ |
| --- | --- | --- |
| Gulf of Mexico (A) | 130.000 | 0.2 |
| Brasil (B) | 20.000 | 0.1 |
| Maine (C) | 60.000 | 0.5 |
| North Atlantic (D) | 60.000 | 0.9 |
| Norway (E) | 40.000 | 0.8 |
| Bay of Biscay (F) | 100.000 | 0.6 |
| Eastern Atlantic (G) | 50.000 | 0.3 |
| Mediterranean sea (H) | 200.000 | 0.2 |

**Table 3.** Seasonal migration phenology of bluefin tuna in the north Atlantic Ocean

| Years | Region | Timing of presence in northern feeding areas | References | Notes |
| --- | --- | --- | --- | --- |
| 1999-2004 | West Atlantic (GOM-Newfoundland); Bay of Biscay | July-September | (Block et al, 2005) | Data Storage Tags (DSTs) |
| 1981-2005 | Bay of Biscay | Day 180-230 (approx.) | (Dufour et al, 2010) | Timing of immigration to region based on commercial CPUE data |
| 2005-2009 | NWAtlantic (Maryland to Cape Cod) | summer months | (Galuardi and Lutcavage, 2012) | DSTs applied to juveniles |
| 1950s-1970s | Norwegian Sea | July-September | (Aloncle et al, 1972; Mather et al, 1995) | Based on commercial catch data. |
| 2012 | East Greenland (Denmark Strait) | August-September | (MacKenzie et al, 2014) | Based on bycatch in commercial mackerel fisheries |
| 1996-2003 | Contl. shelfbreak south of Iceland | August-October (few in November) | (Olafsdottir and Ingimundardottir, 2003) | Based on commercial CPUE data |
| 1923-1931 | Dogger Bank, North Sea | Day 190-290 (approx.) | (Murray, 1932; MacKenzie and Myers, 2007) | Based on at sea observations of schools; similar patterns seen from 1912-1922. |
| 1950s-1970s | North Sea | July-September | (Tiews, 1978) | Based on commercial fisheries |
|  | Southern Gulf of St. Lawrence, Canada | August-October | (Vanderlaan et al, 2014) | Based on commercial CPUE data |
| 1996-2006 | Whole west Atlantic from G. Mexico-Newfoundland, | Seasonal during year | (Walli et al, 2009) | DSTs |

The costs for exploring adjacent habitats, *c_ij_* in Eq. 2,are difficult to set. This is because the term includes several Age class Weight Fertility Growth Survival processes such as traveling between habitats, comparison of habitat qualities and decision making process to select one specific habitat (Bonte et al, 2012). Tuna are efficient swimmers (Dewar and Graham, 1994) and can travel thousand of kilometres within few days (Block et al, 2001). Hence, the cost of traveling and exploring different habitats is not negligible but probably low and most likely dependent on the distance between habitats. Indeed we assume here that the cost for habitat identification and selection is a function of the distance between habitats and approximate it as:

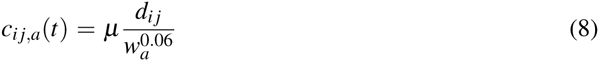

where *d_ij_* is the distance in kilometres between habitat *i* and *j*, while *w*_*a*_ is the age specific average weight, which we assume is proportional to the individual swimming speed. We consider migrations performed at an optimal velocity and it can be shown (Appendix A) that for tunas the swimming speed scales as 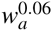 (Ware, 1978). We set the range of *µ* = 5 − 150 to analyse migration game under different habitat selection costs, and we test the sensitivity of our results to this parameter.

The demographic rates in the Leslie matrix (survival *q*_*a*_, fertility *r*_*a*_, growth *g*_*a*_; Eq. 6) are given in Table 1. Fertility coefficients, *r*_*a*_, are non zero only in the spawning areas: Gulf of Mexico (A) and Mediterranean (H). In our definition of the Leslie matrix (Eq. 6) we assume that older individuals have higher fertility proportional to some spawning intensity *s*. In the model we test how the results are affected by different values of *s* (Appendix A).

## 4 Results

### 4.1 Tuna migrations in a stable environment

We first run the model using only the demographic processes, without migration or environmental variability, and set the total bluefin tuna biomass (*M* = 330 *kton*). The simulation converges towards a stable age-structure distribution in the spawning areas (Gulf of Mexico and Mediterranean) and zero biomass otherwise. This is the initial condition used in all the subsequent simulations.

From this initial distribution, we simulate the migration game in the case of a stable environment with no seasonality (*K*_*i*_ constant, Table 2). We assume no demographic changes in the tuna population structure (i.e., the Leslie matrix is the identity matrix) but consider different costs in the habitat selection process. Under such assumptions the migration game on the network converges towards a stable distributional equilibrium (Figure 4).

**Fig. 4.**
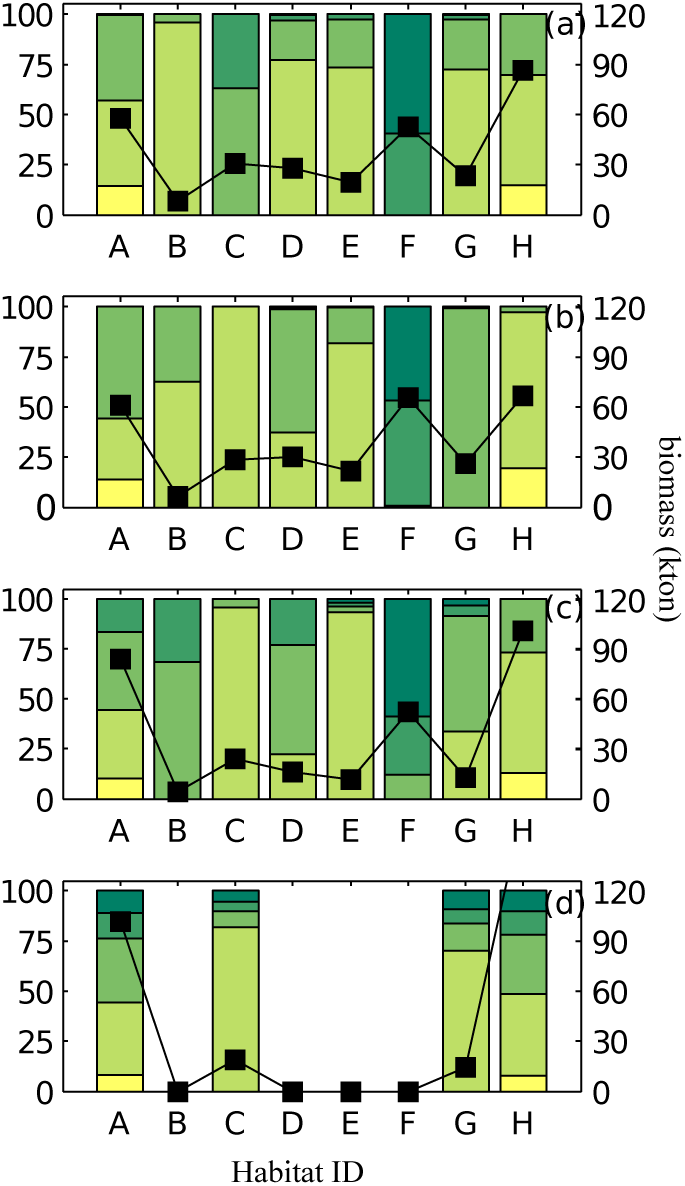
Total biomass (lines) and population structure (bars) in the bluefin tuna habitats as resulting from the equilibrium of the migration game when no demography or seasonal changes are considered in the model. The total biomass of the population is 330 *kt on* while habitat selection costs are (a) very low *µ* = 20, (b) low *µ* = 50, (c) medium *µ* = 100 and (d) high *µ* = 150. Different colours (yellow to dark green) for the 5 different age classes (Young to Old).

At very low costs (*µ* = 20) most of the biomass is aggregated in the spawning areas (*M*_*A*_ = 61 *kton* and *M*_*H*_ = 89 *kton*) and in the Bay of Biscay (*M*_*F*_ = 47 *kton*) while the sum of all the other habitats accounts for ≈ 35% of the total biomass (Figure 4a). In this case largest tuna are on both sides of the Atlantic and mainly in habitats *C* and *F* with Mediterranean and Gulf of Mexico showing the most structured population distribution. The youngest class (yellow color) is present only in the spawning areas and do not migrate in other habitats.

Increasing the habitat selection costs (*µ* = 50, Figure 4b) has no major effects on the biomass distribution. The distributions of age classes are also similar to the previous case with relative changes only in habitats *G* and *C*. With a further increase of the cost (*µ* = 100, Figure 4c) the population tends to accumulate in the spawning areas while the most distant habitats tend to become unoccupied. At very high cost (*µ* = 150, Figure 4c) only few habitats are populated and the majority of tuna biomass is in the spawning area (*M*_*A*_ = 102 *kton*, *M*_*H*_ = 172 *kton*).

### 4.2 Migration game in a seasonal environment

We simulate the habitat selection process under changing carrying capacity (*K*_*i*_) and accounting for tuna population demography (*L*(*t*), Eq. 6).

Seasonal fluctuations in the tuna biomass are evident in all patches with the weakest seasonality in the spawning habitats (Figure 5). The habitat in Norway (Figure 5*E*) and Brazil (Figure *5B*) are occupied by the larger/older classes, but have the lowest of the biomasses. The age-structure in each habitat changes less than the variability in total biomass,but throughout the season significant changes in the age-structure can occur in the eastern Atlantic and Maine (Figure *5C*,*G*). Interestingly the peaks in biomass in the spawning areas are in April - May while in the feeding areas are in July - August and late October in Brazil (Figure 5) as it is commonly reported (Table 4.2).

**Fig. 5.**
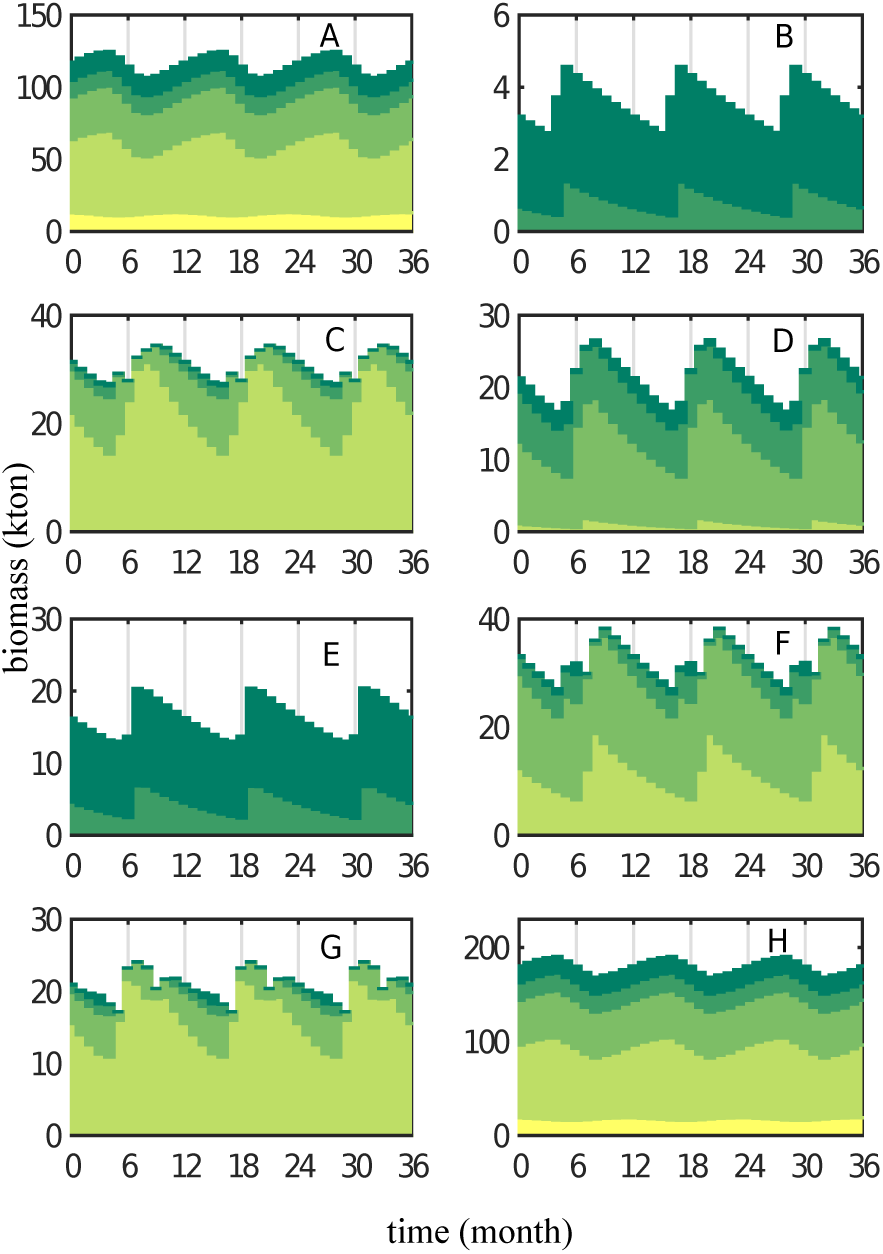
Time series of the biomass per age in the 8 habitats (*A*-*H*) at given habitat selection cost (*µ* = 100) and spawning parameter (*s* = 30). Different colours (yellow to dark green) for the 5 different age classes (Young to Old).

The intensity of migration on the habitat network depends on the cost of the habitat selection process and the spawning intensity of the species (Figure 6). When costs are low and spawning intensity high (Figure 6a) the population distributes in all available habitats and all migratory routes are used with the exception of the transatlantic route C -G. The age-structure is different in each area and highly diversified in the spawning area and in the central Atlantic. When the spawning intensity is reduced (Figure 6b) the total global biomass also decreases and some of the routes are used less frequently. In particular the connections between Brazil and the western Atlantic are much weaker but the transatlantic connections (A -G and C -G) have higher migration flows. This is mainly driven by the very low biomass living in the habitat in Brazil (Figure 6b). At higher habitat selection costs (Figure 6c,d) the direct transatlantic routes connecting habitats A and C to G break down and generally there are low migration rates between habitats. Moreover, only larger individuals appear to exploit the farthest habitats B and E. Further increases of the costs, results in the majority of the population staying in the spawning habitats. In these configurations habitats such as Brazil and Norway have a very low biomass or are completely unoccupied. In the case of high cost and low spawning intensity the migration strategy is only selected by larger individuals while the majority of the population will not distribute outside the spawning grounds. Most of these patterns are confirmed also when a more extensive sensitivity analyses is performed (Appendix A.3).

**Fig. 6.**
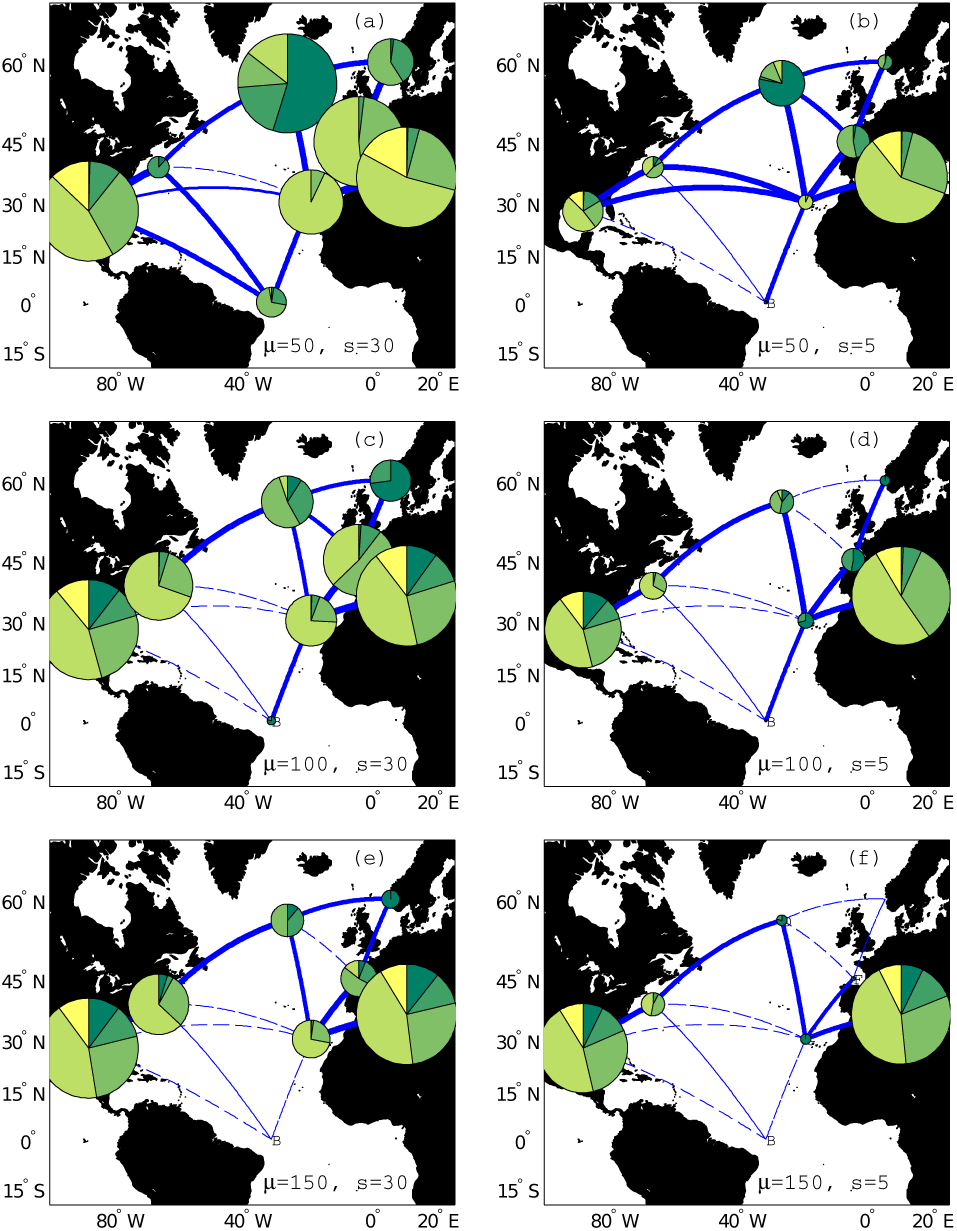
Map of the network structure at different spawning intensity (*s*) and habitat selection cost *µ.* Simulations are run for 10 years under seasonal effects and the distribution in December is shown. Habitats are showed with circle proportional to the total tuna biomass and colour for different age (between yellow and dark green from Young to Old age, respectively). Lines connecting the habitats show integral biomass flux during the entire simulation and are thicker for larger fluxes, while dashed lines are used when no flow is simulated along the path. (a) *µ* = 50 and *s* = 30 and a total global biomass *M* = 496 *kt on*, (b) *µ* = 50 and *s* = 5 and *M* = 128 *kt on*, (c) *µ* = 100 and *s* = 30 and *M* = 424 *kt on*, (d) µ = 100 and *s* = 5 and *M* = 166 *kt on*, (e) µ = 150 and *s* = 30 and *M* = 374 *kt on*, (f) *µ* = 150 and *s* = 5 and *M* = 174 *kt on*.

## 5 Discussions

### 5.1 Migration modelling

We introduce a model based on game theory to simulate habitat selection processes in migratory populations. The model is developed to describe migration dynamics in age structured fish populations and it is applied to study the seasonal migration of the Atlantic bluefin tuna. The model explicitly represents habitat connectivity as a network with several patches connected by links.

The results show how changes in the resource level, population demography and cost of migrations can alter the migration equilibrium of the game and then results in different population distributions in the habitats. We further show that only some subset of the available links on the network are effectively selected as migratory pathways while many other routes are not utilised. This allows us to identify emerging migration routes in fish populations and to compare the predictions with observed migration behaviour.

A fundamental assumption in the model is that migration is described as a multiple equilibrium process between different habitats. In the migration game, each individual in a given habitat can - in a single time step - move only to the neighbour habitats, i.e., those that are locally connected to the one where the individual is living. The migration occurs when there is an advantage to move, which in the model is described as a positive reward function. Since this function is negatively density dependent, its value is affected by the strategies of other individuals in the populations and it is also affected by the cost of assessing and commuting between different habitats. At each time step the population tends to reach a local equilibrium by trying to equalise the local reward functions. In some cases this equilibrium cannot be reached because there are not enough individuals living in a given habitat that can migrate towards connected habitats with a higher payoffs.

The assumption of describing habitat connectivity by discrete network structure is a generalisation of migration models assuming movements between all pairs of habitats (a fully connected graph). The network approach easily captures the existence of geographical, bioenergetic or life history constrains, which often break potential migration routes (Henningsson and Alerstam, 2005; Alerstam et al, 2003; Alerstam, 2001). The model is also flexible enough to allow the effects of ocean currents, temperature variability, or other environmental changes to be represented using different costs on each link. Indeed, the cost of migration between two habitats can affect the reward function and then can modify the migration equilibrium on the network; a mechanism which is in agreement with the hypotheses that changes inmigration routes can be driven by climate change (Walther et al, 2002; Rijnsdorp et al, 2009; Doney et al, 2012).

The ability to equalise the local reward functions and reach an equilibrium is consistent with the ideal free distribution theory (Fretwell and Lucas, 1969). We show that at each time step individuals in the population distribute according to a local IFD among connected habitats and, in case of stable environment with no demographic effects, the local equilibrium converges, in several steps, towards a global IFD on the network (Pan and Nagurney, 1994; Cressman and Křivan, 2006). The behaviour dynamic we use in the model describes when and how individuals update their strategies over time. This is known as revision protocol in game theory (Sandholm, 2010) and is based on two assumptions: myopia and inertia. A myopic behaviour means that individuals assess their strategy based on local information on costs and payoff opportunities, without incorporating knowledge on future expectations and behaviours. Inertia in behaviour considers that individuals do not update their strategy continuously but instead re-evaluate their decision sporadically, mainly because very often the environment in which they live provides a multiplicity of concerns to be solved rather than a single-minded focus on one strategy (Sandholm, 2010). We think that the discrete form of time and space in our model describes naturally myopic and inertial processes which are likely to occur in fish populations. Moreover, compared to a continuous model, the discrete representation of the space with the network of habitats appears to be more consistent with the idea of migration corridors, habitat hot spots and migration stopovers which are typically found in many species (Rose, 1993; Hunter et al, 2003).

### 5.2 Environmental and demographic effects

Seasonality in the resource distribution and competition for resources are both important mechanisms for the selection of migration as a behavioural trait. Indeed, in weak seasonal environments and under low competition conditions, residency is the behavioural strategy that is selected (Shaw and Couzin, 2013). Our results support those findings and show that in case of a stable environment the distribution of the populations between heterogeneous habitat converges to the IFD. At the IFD there is no net dispersal between habitats unless demographic effects are present. Indeed changes in population structure can elicit changes in the reward functions then allowing migrations to emerge. The coupling between demographic and environmental effects are critical for the results. Generally, if dispersal dynamics are much faster than the changes in the habitats payoff, the IFD can still be reached by migratory species. But if migrations and environmental variability have similar time scales, additional assumptions need to be made on dispersal (Mobæk et al, 2009;McLoughlin et al, 2010). In our model we assume that the population can reach a local IFD condition, which translates into assuming a fast behavioural response to explore neighbour habitats. The time step used to integrate the discrete model sets the time scales for behavioural response and environmental variability. An alternative approach would describe demographic and environmental dynamics as continuous processes both affecting the migratory behaviour with feedback on the population dynamics. In such cases however we would have a revision protocol lacking the inertia of the decision process which we think is common in most natural populations.

The presented model does not explicitly account for the feedbacks between migration dynamics and demographic processes. Indeed, in habitats with higher payoff one could expect individuals to grow faster than those in lower payoff habitat. To capture the impacts of a large payoff on individual growth, one would likely require an individual-based approach to store information about memory and history of the single individuals.

### 5.3 Relevance to the ecology of the Atlantic bluefin tuna

As we have seen before, Atlantic bluefin tunas have a wide distribution in the Atlantic Ocean from tropical to sub-polar areas. Migration has likely evolved to allow migrants to benefit from the highly productive environment at higher latitude while reducing competition by moving towards less productive environments. Being excellent swimmers Bluefin tuna can potentially be present in all parts of the Atlantic. Nonetheless, several evidences suggest that the species distributes within several hotspots areas, where tunas are present all year round, while their abundance outside those areas are minimal. Moreover, the same individual can visit these hotspots several times during the feeding period before going back to spawning areas for reproduction. Those patterns in distribution and migration behaviour are in part captured by the habitat network approach used in our model, with a series of hotspot areas connected by a range of migratory pathways. Moreover, the model appears to describe reasonably well the peaks in distribution in the different areas. For example in the spawning areas the maximum abundance is mainly at the beginning of summer and precedes the peaks in abundances in the feeding areas. Habitats such as Norway or Brazil are visited only by the larger individuals (200 – 500*Kg*) and are very sensible to changes in fishing pressure or cost of migrations (Fromentin, 2009; Safina and Klinger, 2008).

Thus our modelling approach allows representing in a quite realistic way the spatial population dynamics of the Atlantic bluefin tuna including simulating the disappearance of previous feeding habitats and changes in migration routes.

The modelled estimate of the timing of appearance at summer feeding areas is similar to the migration phenology to many of these areas observed in nature (Table 4.2). In addition, the size composition of the modelled populations arriving in several of these areas compares favourably with the size composition of bluefin tuna observed and / or caught in such regions.

For example, modelled size distributions for Brazil and Norway are centred at large (≈ 200 *cm*) sizes; catch data from these areas (Mather et al, 1995) shows that most bluefin captured in fisheries in these areas were generally > 150 – 200 *cm*, thus similar to modelled estimates.

Our modelling approach is potentially a useful framework for investigating how exploitation and environmental variability including climate change could affect the largescale migratory behaviour and spatial distribution of Bluefin tuna and its phenology. For example, environmentally driven changes in regional productivity and carrying capacity would affect habitat suitability, migration costs (e.g., due to temperature changes) and migratory rewards after arriving at destinations. These changes could lead to reductions of utilisation of some habitats and stronger preferences for habitats in other regions, thereby potentially influencing fishery opportunities and costs for different nations. Such changes may already be underway because migration phenology for the Bay of Biscay is linked to large-scale climate conditions (e.g., the North Atlantic Oscillation, NAO) that affect sea temperatures (Dufour et al, 2010) and bluefin tuna have recently been observed in east Greenland where they have not previously been observed (MacKenzie et al, 2014). Moreover our modelling framework, if coupled to integrated oceanographic biogeochemical models (Dragon et al, 2015), could also potentially be used to derive new insights on the relative roles of oceanographic variability and exploitation leading to past major changes in bluefin tuna distributions and fisheries such as those off Brazil, Norwegian-North Sea and south of Iceland. Recent advances in group behaviour and information sharing/transfer between individuals within groups also show how habitat choice can be influenced by the knowledge content or migratory experience of individuals within groups, and how group behaviour (e.g., migration to particular habitats) can be driven by a subset of informed individuals (De Luca et al, 2014).

A future challenge for migratory behaviour modelling is therefore to develop ways to integrate individual-level and group dynamics in migration game modelling frameworks such as that developed here. Given that bluefin tuna is such a highly migratory species, and migrates across ocean zoning boundaries of several jurisdictions, and also across stock management boundaries, migration models that quantify rates and timing of exchanges among areas could potentially have practical application in fishery management and conservation. The migratory behaviour of this species is complex. Our modelling approach, although moderately complex, is based on some simplified considerations of population dynamics, regionally-dependent ecosystem carrying capacities and bioenergetics of energy intake and utilisation, and is a step towards process-oriented migration and distribution models. Further advances in process knowledge and implementation are needed, and if implemented, could support management and conservation decision-making for this species.

## Acknowledgements

The Authors wish to thank the participants to the conference ’Dispersal and competition of populations and communities in spatially heterogeneous environments’, Lausanne, Switzerland, 4-8 August 2014 for inspiring some of this work. PM received support from Otto Mønsted Fond and was supported by the European Union Seventh Framework Programme project EURO-BASIN (ENV.2010.2.2.1-1) under grant agreement nr. 264933. This work was partly conducted while VK was a Sabbatical Fellow at the Mathematical Biosciences Institute, an Institute sponsored by the National Science Foundation under grant DMS 0931642. Support provided by the Institute of Entomology (RVO:60077344) is acknowledged.

## A Model calibration for Atlantic bluefin tuna

### A.1 Migration costs

The time needed to migrate between two habitats regulates the cost of migration in fish population since the energy consumed will be higher the longer is the migration time. The power rate consumed while swimming at an optimal speed (*P*) is:

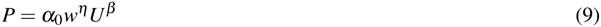

where *w* is the weight of the fish, while the allometric constants *α*_0_ and η have been estimated for fish swimming at high Reynolds number (Ware, 1978, Table 4).

**Table 4.** Scaling of physiological rates with size and parameter values for tuna from: [1] Overholtz (2006) [2] Ware (1978) [3] Dewar and Graham (1994) [4] Block and Stevens (2001)

| Parameter | Symbol | Value | Ref. |
| --- | --- | --- | --- |
| Max. Energy intake | $I(w)$ | $cw^\phi$ | – |
| Standard metabolic rate | $M(w)$ | $\alpha_1 w^\gamma$ | – |
| Swimming power | $P(w)$ | $\alpha_0 w^\eta U^\beta$ | – |
| Constant for energy intake | $c$ | $1 \cdot 10^{-2}$ | [1] |
| Exponent for energy intake | $\phi$ | 0.8 | [1] |
| Constant for power cost | $\alpha_0$ | $1.8 \cdot 10^{-8}$ | [2] |
| Exponent for power cost | $\eta$ | 0.47 | [2] |
| Exponent for power cost | $\beta$ | $1.4 < \beta < 2.8$<br>($\beta = 2$) | [3] |
| Constant for metabolic cost | $\alpha_1$ | $3.76 \cdot 10^{-4}$ | [4] |
| Exponent for metabolic cost | $\gamma$ | 0.6 | [4] |

We can fairly assume that migrations is performed at an optimal swimming speed (*U*^*^) at which the total energy expenditure per unit distance is minimised. Using an allometric function for the metabolic costs *M* = α_1_*w*^γ^, a general form of *U*^*^ can be derived by an optimisation procedure relating the swimming cost to the total cost of moving (metabolic cost plus power output):

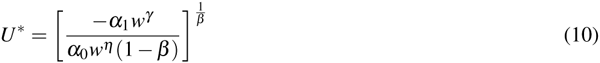

where α_1_ and γ are allometric constants for fish metabolism (Table 4). This results in an allometric scaling for the optimal swimming speed as:

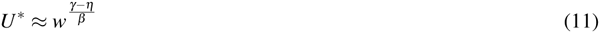

In tuna the exponent *β* has been found to range between 1.4 < *β* < 2.8 (Dewar and Graham, 1994) and we assume *β* = 2.1, which provides swimming speeds in the range reported for several tuna species (1.2 − 2.4*BLs*^−1^) (Block and Stevens, 2001).

Thus we obtain a scaling *U*^*^≈ *w*^0.06^.

### A.2 Demography

Large uncertainties exist on the definition of demographic parameters for the bluefin tuna population (Simon et al, 2012). In our model, the young-of-the-year stage (0 -1 years) is excluding egg phases and does not have reproductive potential while at juvenile stage (1 -5 years) a small fraction is mature for reproduction. The reproductive maturity increases up to 50% at the adult stage (5 -10 years) while mature (10 -20) and old (20 -35) stages are fully reproductive but the latter has a lower survival rate. Those rates are consistent with observed maturity at age data for western and eastern atlantic bluefin tuna (SCRS, 2012) and are used to define the values of *r*_*k*_. Moreover, the value survival (*q*) and growth (*g*) values used in the Leslie matrix are consistent with reported values for the yearly mortality rates (SCRS, 2012) and provide a realistic bluefin tuna age-structure (Fig. 7) with a maximum population growth rate (0.15) that is in the range of previous estimates (Simon et al, 2012). Finally, we constrain the global bluefin tuna population using a given total carrying capacity *K*_*t*_ and assume a density dependent function on the spawning factor *s*.

**Fig. 7.**
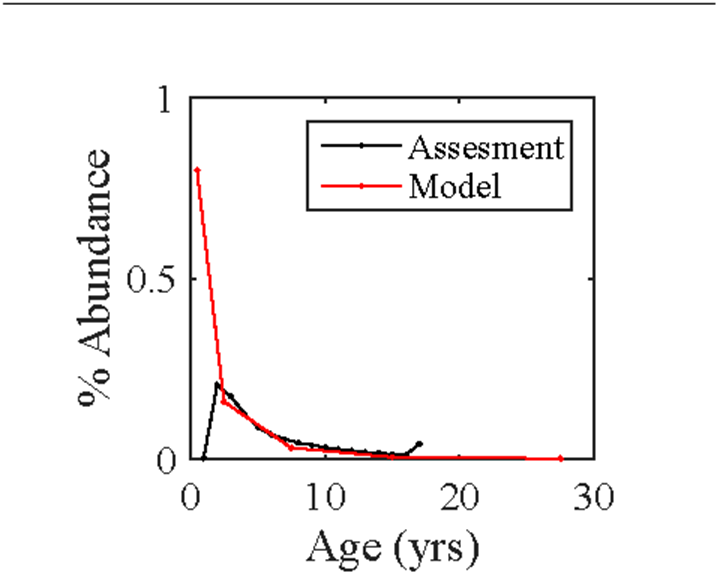
Age structure data from the ICCAT assessment group (black) on bluefin tuna and from the model using the Leslie matrix estimates (red).

### A.3 Extended Sensitivity analyses

At low spawning intensity and high migration costs (Figure 8g) only the spawning areas are occupied. Decreasing habitat selection costs allows tuna to migrate in adjacent feeding areas (G and C) but reduce the total biomass and increase fluctuations in the migration behaviour (Figure 8a,d). On the other hand, at high spawning and low migration costs (Figure 8a,b) the biomass reaches the total carrying capacity over few months, and all habitats are occupied although at different levels of biomass.

**Fig. 8.**
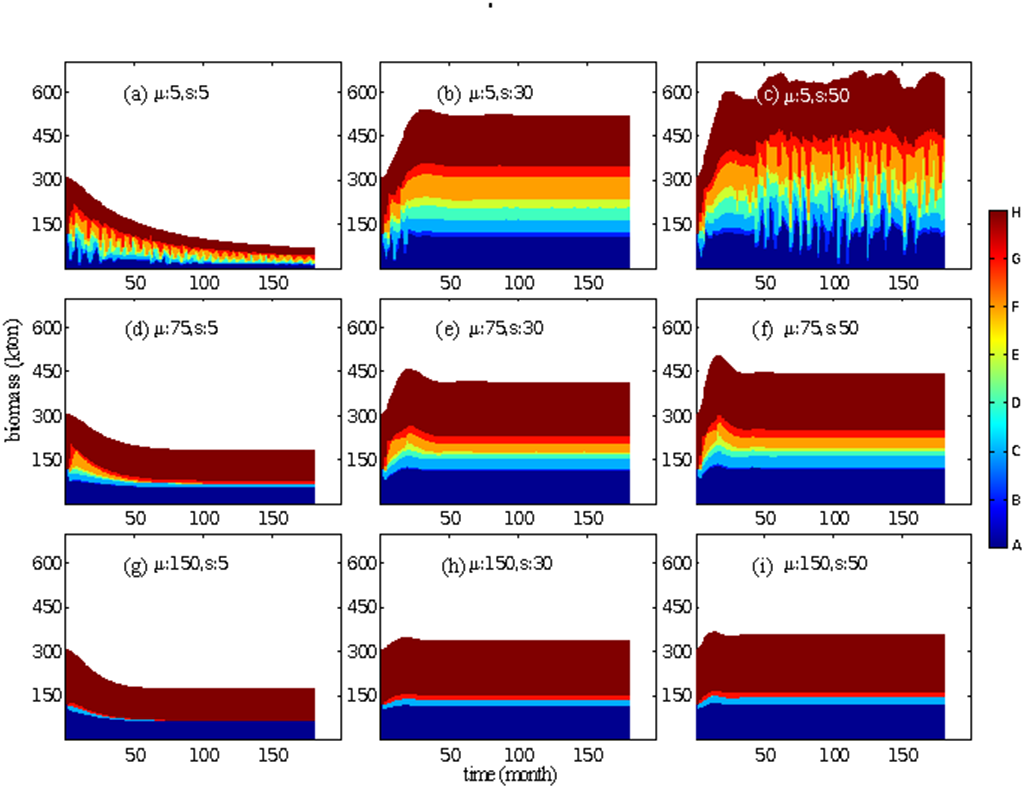
Sensitivity of the population structure and total biomass in different habitats in case of no seasonality and zero fishing under different spawning intensity s and habitat selection cost *µ.*

